# Species-specific stomatal ABA responses in juvenile ferns grown from spores

**DOI:** 10.1101/2023.03.09.531841

**Authors:** Tana Wuyun, Ülo Niinemets, Hanna Hõrak

**Affiliations:** Institute of Agricultural and Environmental Sciences, Estonian University of Life Sciences, Kreutzwaldi 1, 51006 Tartu, Estonia; Institute of Technology, University of Tartu, Nooruse 1, 50411 Tartu, Estonia

**Keywords:** ferns, ABA response, stomatal physiology

## Abstract

Adjustable stomatal pores in leaves control the balance between CO_2_ entry for photosynthesis and water loss via transpiration. The drought and low humidity-induced phytohormone abscisic acid (ABA) is the major regulator of active stomatal closure responses in angiosperms. Whether the ABA signalling pathway for stomatal closure functions similarly in older land plant groups, such as lycophytes and ferns, is still unclear: some studies find no stomatal ABA response in ferns, others find that ABA response is present or triggered by specific environmental conditions. Here we analysed steady-state gas-exchange, stomatal density and stomatal response to exogenously applied ABA in nine fern species grown from spores under controlled growth conditions. We find that ABA responses in ferns are species-specific: stomata in four out of nine species closed in response to ABA. The ABA-sensitive species mostly had slow responses of low magnitude, suggesting reduced ABA-sensitivity of ABA signalling pathway in ferns. Species with larger stomatal conductance tended to close stomata in response to ABA, whereas a relatively strong response of ~35% was also found in *Cyrtomium falcatum*, a fern with low stomatal conductance. Our results show that ferns constitute a diverse group with varying degree of stomatal ABA-sensitivity. Further characterisation of ABA signalling pathway components in diverse fern species is needed to understand the genetic basis for the variable ABA-sensitivity in ferns.

## Main text

Stomatal pores in leaves control CO_2_ entry for photosynthesis and water loss via transpiration. Stomata open in response to light, low CO_2_ levels and high air humidity, and close in response to darkness, elevated CO_2_ concentration, low air and soil humidity. Stomatal closure responses are largely mediated by the stress hormone abscisic acid (ABA) that is produced in response to drought and reduced air humidity (Xiong & Zhu, 2003; McAdam *et al*., 2016b). ABA is perceived by ABA receptors from the PYR/PYL family that form a complex with PP2C phosphatases and suppress their activity (Ma *et al*., 2009; Park *et al*., 2009; Melcher *et al*., 2009). Suppression of PP2C activity allows the activation the OST1 kinase that phosphorylates and activates the SLAC1 anion channel, triggering ABA-induced stomatal closure (Vahisalu *et al*., 2008, 2010; Negi *et al*., 2008; Fujii *et al*., 2009; Geiger *et al*., 2009; Lee *et al*., 2009). How the molecular mechanisms that coordinate stomatal responses to environmental change and ABA have evolved has intrigued stomatal biologists over the last decade.

Lycophytes and ferns are one of the oldest plant groups with stomata (Clark *et al*., 2022). Ferns have been proposed to completely lack active ABA-induced stomatal closure and to close stomata via solely hydropassive mechanisms (Brodribb *et al*., 2009; Brodribb & McAdam, 2011; McAdam & Brodribb, 2012; Cardoso *et al*., 2019). At the same time, ABA-induced stomatal closure has been documented in several fern and lycophyte species (Ruszala *et al*., 2011; Cai *et al*., 2017; Hõrak *et al*., 2017; Tai-Chung Wu, 2020). Several studies have also indicated that fern stomatal responses are species-specific and vary with growth conditions (Hõrak *et al*., 2017; Plackett *et al*., 2021). The species-specificity may be explained by different numbers of ABA receptors and varying degree of their ABA-sensitivity in different species (Sun *et al*., 2019), and different degree of activation of the SLAC1 anion channel via OST1 homologs of different fern or lycophyte species (Ruszala *et al*., 2011; McAdam *et al*., 2016a). The conditionality of fern stomatal responses may result from different lifetime growth conditions leading to e.g. differences in the endogenous ABA levels (McAdam & Brodribb, 2012; Cardoso *et al*., 2019). Previous studies on fern stomatal physiology have often focused on rhizome-grown mature plants. Changes in environmental conditions experienced throughout the lifetime may have resulted in adaptations to local growth environment in such plants, potentially introducing legacy effects that affect their response to external cues. Growth conditions affected ABA-responsiveness in the ferns *Athyrium filix-femina* and *Dryopteris filix-mas* (Hõrak *et al*., 2017), and priming of ABA responses was found in *Ceratopteris richardii*, where stomata became ABA-responsive after pre-treatment with ABA (Plackett *et al*., 2021). Here, we aimed to minimize priming and life history effects via analysis of nine fern species grown from spores under controlled growth conditions: *Adiantum capillus-veneris, Adiantum polyphyllum, Cheilantes distans*, *Cheilantes farinosa*, *Cheilantes lanosa*, *Asplenium scolopendrium*, *Blechnum occidentale*, *Cyrtomium falcatum*, and *Dryopteris sieboldii*. We then analysed steady-state gas-exchange, stomatal density and ABA-responsiveness in the juvenile leaves of these ferns to resolve the physiological and anatomical controls on fern stomatal responsiveness to ABA.

We germinated the spores under humid conditions and achieved gametophytes (Fig. 1a) that successfully fertilised and gave rise to sporophytes that could be analysed in the leaf chamber of a gas-exchange system (Fig. 1b; see also details in Materials and Methods). In line with previous studies on mature ferns (Hõrak *et al*., 2017; Xiong & Flexas, 2020; Cândido-Sobrinho *et al*., 2022), stomatal conductance (*g*_s_) and net photosynthesis rate (*A*_net_) in juvenile ferns were generally low (Fig. 1c, d,). *g*_s_ varied 7-fold between the studied species, ranging from 20 to 150 mmol m^-2^ s^-1^ (Fig. 1c), whereas variation in *A*_net_ was even larger, ranging from 0.5 to 5 μmol m^-2^ s^-1^ (Fig. 1d). Ferns from the *Cheilantes* genus had the highest *g*_s_ values in our study. There was a strong positive relationship between *g*_s_ and *A*_net_ (Fig. 1e, *R*^2^=0.68, *P*=0.004), similar to mature ferns (Kübarsepp *et al*., 2020) and across angiosperms (Wright *et al*., 2004).

**Figure 1.**
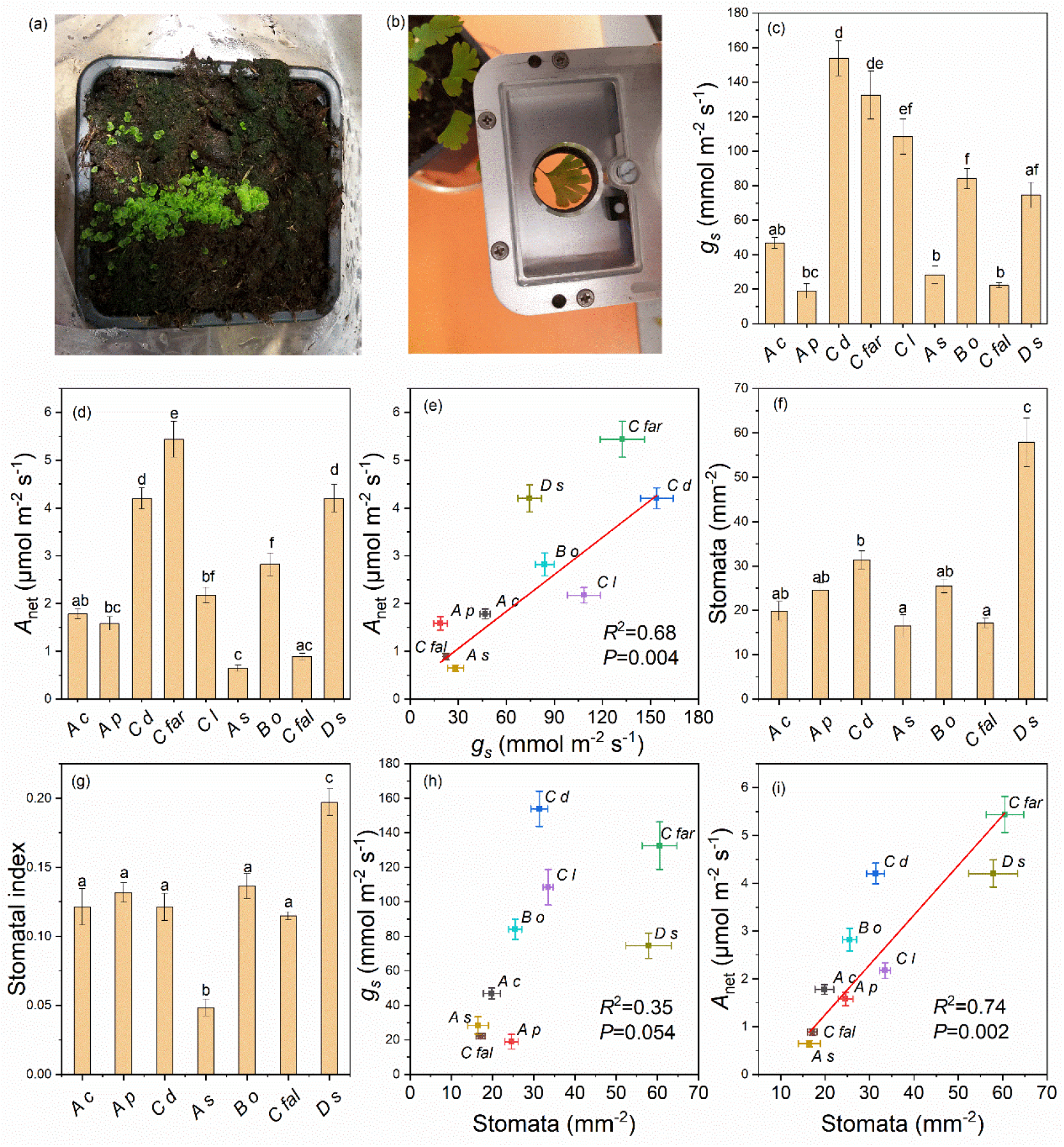
Steady-state gas-exchange traits of juvenile ferns. Representative image of *C. falcatum* gametophytes (a), representative image of *A. capillus-veneris* in leaf gas-exchange chamber (b), steady-state pre-treatment stomatal conductance to water vapor (*g*_s_, c), steady-state pre-treatment net assimilation rate (*A*_net_, d), relationship between *g*_s_ and *A*_net_ (e), stomatal density (f), stomatal index (g), relationship between stomatal density and *g*_s_ (h), and relationship between stomatal density and *A*_net_ (i). Species codes as: *A c – A*. *capillus-veneris*, *A p – A*. *polyphyllum*, *C d – C*. *distans*, *C far* – *C. farinosa*, *C l* – *C. lanosa*, *A s* – *A. scolopendrium*, *B o* – *B. occidentale*, *C fal* – *C. falcatum*, *D s* – *D. sieboldii*. Mean ± SEM is shown in panels c-i, *n* = 5-12 plants. One-way ANOVA with Tukey *post hoc* test was used in panels c, d, f and g, different letters show statistically significantly different groups. Linear regression was used in panels e, h, and i.

Ferns are known to have large and sparse stomata (Lima *et al*., 2019; Kübarsepp *et al*., 2020; Xiong & Flexas, 2020). In line with previous studies, we found low stomatal density in juvenile leaves of the studied species, with most values ~20 stomata mm^-2^. *Dryopteris sieboldii* had the highest stomatal density of about 50 stomata mm^-2^ (Fig. 1f). In most species, stomatal index (the proportion of stomata from all epidermal cells) was ~0.12 with a lower value observed in *A. scolopendrium* and a higher in *D. sieboldii* (Fig. 1g). Compared to typical stomatal density and index values in angiosperms, ferns strike out has having few large stomata (Kübarsepp *et al*., 2020)). This might contribute to the sluggishness of their stomatal responses, as stomatal size and speed are negatively correlated (Drake *et al*., 2013; Kübarsepp *et al*., 2020). We found no significant relationship between stomatal density and *g*_s_ (Fig. 1h), but in our sample, there was a linear positive relationship between stomatal density and *A*_net_ (*R*^2^=0.74, *P*=0.002, Fig 1i). These data suggest that in our experiments, *A*_net_ was indirectly affected by stomatal density, potentially via simultaneous adjustment of stomatal density and anatomical factors determining photosynthetic capacity, such as leaf thickness and mesophyll structure.

To test if juvenile ferns grown from spores under controlled conditions can respond to ABA, we spray-applied 10 μM ABA or mock solution (see details in Materials and Methods) and followed stomatal conductance in treated leaves for 1.5 hours after the treatment (Fig. 2a-i, Fig. S1). The low stomatal conductance of ferns was in some cases accompanied by large variation and fluctuations in *g*_s_ (e.g. Fig. 2b, f), and in two *Cheilantes* species mock-treatment also affected *g*_s_ (Fig. 2c, e). However, it was possible to clearly differentiate ABA-responsive and unresponsive fern species in our sample. External ABA triggered significant stomatal closure compared to mock-treatment in *C. distans, C. lanosa, B. occidentale* and *C. falcatum* (Fig. 2c, e, g, h, j), whereas no difference between mock- and ABA-treated leaves was observed in *A. capillus-veneris, A. polyphyllum, C. farinosa, A. scolopendrium*, and *D. sieboldii* (Fig. 2a, b, d, f, i, j). In ABA-responsive ferns, the responses were slow and relatively low in magnitude (Fig. 2c, e, g, h, j). This might explain why no ABA responses were previously found in shorter experiments (Cândido-Sobrinho *et al*., 2022).

**Figure 2.**
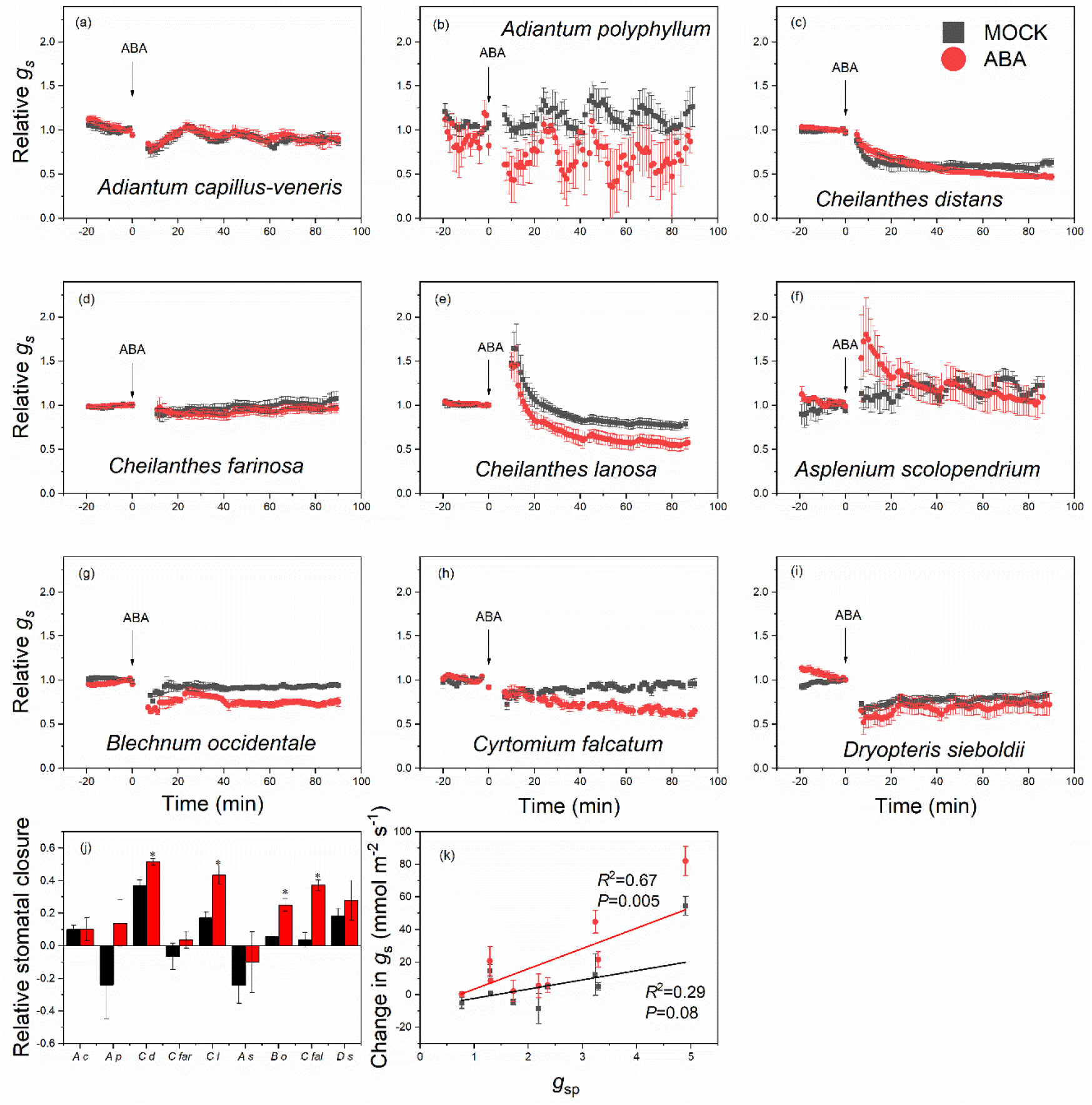
Stomatal ABA responses in juvenile ferns. Stomatal response in relative units (relative to conductance at time point 0) to treatment with 10 μM ABA or mock-solution in (a) *A. capillus-veneris*, (b) *A. polyphyllum*, (c) *C. distans*, (d) *C. farinosa*, (e) *C. lanosa*, (f) *A. scolopendrium*, (g) *B. occidentale*, (h) *C. falcatum*, and (i) *D. sieboldii*, and (j) change in *g*_s_ in relative units between 0 and 90 minutes after treatment, (k) relationship between *g*_s_ per million stomata (*g*_sp_) and absolute change in *g*_s_. In a-i, ABA or mock treatment were applied at time point 0. Mean ± SEM is shown in all panels, *n* = 4-5 plants. Multiple t-tests with Bonferroni correction were used in panel j, asterisks show statistically significant effect of ABA treatment. Linear regressions were used in panel k.

Ferns with relatively high initial *g*_s_ tended to close stomata in response to ABA (Fig. 1a, 2c, e, g, j), potentially suggesting that higher doses of ABA received through open stomata may account for the differences in ABA response between species. To account for differences in stomatal densities between species, we calculated normalized stomatal conductance (*g*_s_ per million stomata, *g*_sp_), and analysed its relationship with ABA-induced stomatal closure. There was a significant positive linear relationship between *g*_sp_ and ABA-induced change in *g*_s_ (*R*^2^=0.65, *P*=0.005, Fig 2k) that was absent in mock-treated plants (*R*^2^=0.29, *P*=0.08). Thus, it is possible that different ABA responses between fern species may be to some extent explained by different ABA doses that reach guard cells in ferns with different stomatal apertures and densities. Nevertheless, in our study, the fern *C. falcatum* that had one of the lowest *g*s before treatment (Fig. 1c) and hence potentially low ABA uptake, showed a clear and relatively strong (~35%) ABA-specific stomatal closure, suggesting high ABA-sensitivity of the ABA signalling pathway in this species.

Even the species that closed stomata in response to ABA had a slow response with variable, but relatively small magnitude, as documented before for mature ferns (Hõrak *et al*., 2017; Plackett *et al*., 2021). Thus, although some fern species can respond to ABA, the mechanisms underlying these responses, or their ABA-sensitivity, are likely at least partly different from angiosperms. The reduced ABA-sensitivity of ferns may be explained by lower ABA-sensitivity of fern ABA receptors (Sun *et al*., 2019). In addition, the anatomical characteristics of fern stomata (large stomata with reduced mechanical advantage of epidermal cells over guard cells (Franks & Farquhar, 2007; Franks, 2013; Xiong & Flexas, 2020)) may contribute to slow stomatal responses in ferns.

Due to the lack of humidification in our growth room (see Materials and Methods), it is possible that the stomata in all studied species were primed for ABA-responsiveness by the relatively dry indoor air as found before for *C. richardii* (Plackett *et al*., 2021). Even under these conditions, five of the nine studied species were completely insensitive to ABA, suggesting that while fern stomatal responses can be triggered conditionally in some species, ABA-responsiveness in ferns is species-specific and absent in many species, in line with previously reported ABA-insensitivity of some ferns (Brodribb & McAdam, 2011; Gong *et al*., 2021). As the ABA signalosome (Sun *et al*., 2020) and its role in stomatal responses (Clark *et al*., 2022) are considered an ancestral state present in the common ancestor of land plants, the absence of ABA responses in some ferns may be explained by secondary loss of ABA-response pathway components, or reduction of ABA-sensitivity of these components. Variation in ABA-sensitivity between ABA receptors is known to occur both within and between species (Sun *et al*., 2019), and it can be expected to translate into different stomatal ABA-sensitivity. Hence, species-specificity of fern stomatal ABA responses is expected, although environmental conditions experienced throughout the lifetime likely affect both anatomical traits and physiology of stomatal responsiveness in ferns (Hõrak *et al*., 2017; Plackett *et al*., 2021).

Stomatal closure responses to elevated CO_2_ have often been discussed together with ABA-responsiveness in ferns, implying similar underlying molecular mechanisms for these responses. Most studies have found that ferns close their stomata in response to elevated CO_2_, although the response is lower in magnitude and slower than in angiosperms (Franks & Britton-Harper, 2016; Hõrak *et al*., 2017; Lima *et al*., 2019; Kübarsepp *et al*., 2020; Tai-Chung Wu, 2020). No response has been observed in a few cases (Brodribb *et al*., 2009; Brodribb & McAdam, 2013). While responses to elevated CO_2_ are indeed partly mediated via the ABA pathway in angiosperms (Merilo *et al*., 2013; Chater *et al*., 2015; Dittrich *et al*., 2019), and thus possibly also in ferns, in the model angiosperm *Arabidopsis thaliana*, CO_2_-induced stomatal closure largely relies on the signalling module involving the HT1 and MPK12 kinases (Hashimoto *et al*., 2006; Hõrak *et al*., 2016; Jakobson *et al*., 2016; Takahashi *et al*., 2022). An HT1 homologue is expressed in the stomata-bearing sporophyte of the moss *Physcomitrium patens* (O’Donoghue *et al*., 2013; Chater *et al*., 2013), but whether its function is conserved from mosses to angiosperms is unclear (Harris *et al*., 2020). Thus, analysis of fern HT1 pathway genes and their functional conservation will be important to understand the evolution of signalling pathways that control plant stomatal responses to changes in CO_2_ concentration. Given the stomata-specific role of the HT1-MPK12 pathway in *A. thaliana*, it can be a good indicator for evolution of stomatal CO_2_-responsiveness that is not confounded by pleiotropic effects that can be expected for ABA, a hormone with diverse roles in plants.

Our study using ferns grown from spores under controlled conditions indicates that fern species-specific stomatal responsiveness is genetically determined and cannot be explained solely by different life history and growth conditions experienced over the plant lifetime. The mounting evidence on the variability of fern stomatal responses to ABA across species suggests that ferns constitute a diverse group in terms of their ABA responses. As sequencing and -omics approaches become widely accessible, it will be possible to characterise the ABA- and CO_2_-response pathways in more fern species to better understand the evolution of stomatal responsiveness. Our study shows that there are ABA-insensitive species, and also that in the ABA-sensitive species, the responsiveness is often stronger in species with overall greater capacity for water loss (higher *g*_s_). Thus, fine-tuning of fern water use can be particularly relevant for species with potentially greater physiological activity.

## Materials and methods

### We included nine fern species in our study

*Adiantum capillus-veneris* (University of Tartu Botanical Garden), *Adiantum polyphyllum* (University of Tartu Botanical Garden), *Cheilantes distans* (University of Tartu Botanical Garden), *Cheilantes farinosa* (University of Tartu Botanical Garden), *Cheilantes lanosa* (University of Tartu Botanical Garden), *Asplenium scolopendrium* (University of Tartu Botanical Garden), *Blechnum occidentale* (New Zealand), *Cyrtomium falcatum* (Estonian University of Life Sciences), and *Dryopteris sieboldii* (University of Tartu Botanical Garden). We grew the plants in a growth room (temperature 23 °C, photoperiod 12 h/12 h, moderate growth light level 50 μmol m^-2^ s^-1^ reflecting the conditions in the forest floor, relative humidity was not controlled) throughout the experiments. We sowed fern spores into 0.29 l (8 x 8 x 7 cm, length x width x height) plastic pots (Teku, Pöppelmann, Germany) with wet soil (commercial garden soil (N:P:K = 10:8:16, Kekkilä Group, Vantaa, Finland), and perlite in a 1:1 volume ratio) and incubated these in transparent plastic bags to preserve humidity and stimulate growth of gametophytes (Fig. 1a) and fertilization. When the first leaves of sporophytes emerged, we removed the plastic bags and transplanted individual sporophytes to separate pots for subsequent growth. After 4-8 juvenile leaves had reached sufficient size to be analysed by a portable gas-exchange system (GFS-3000, Walz GmbH), we carried out measurements of stomatal conductance (*g*_s_) and net photosynthesis (*A*_net_) under the following conditions: 300 μmol m^-2^ s^-1^ light, 70% relative humidity, 400 μmol mol^-1^ CO_2_, cuvette temperature 25 °C. Thereafter, we subjected each measured plant to either mock (0.01% surfactant Silwet L-77 in water) or ABA (10 μM ABA, 0.01% Silwet L-77 in water) treatment. Briefly, we removed the analysed leaf from the leaf cuvette, sprayed it with treatment solution, air-dried the leaf and inserted it back into the cuvette within 1-5 min, taking care to clamp the same part of the leaf into the cuvette as done before the treatment. We followed stomatal conductance of the treated leaf for 1.5 hours to assess stomatal response to ABA. We calculated relative stomatal response curves with respect to pre-treatment *g*_s_ and relative and absolute changes in *g*_s_ during the 90 minutes after treatment (change in *g*_s_ in relative units or in mmol m^-2^ s^-1^ between *g*_s_ at time point 0 and *g*_s_ at 90 min after treatment). To characterise stomatal density and index, we captured dental resin (Xantopren L blue, Kulzer) imprints from lower (abaxial) side of leaves, and then applied nail varnish to derive transparent impressions as in Casson *et al*. (2009). We used a brightfield microscope (OBF 133C832, with digital camera ODC 832, Kern) and MicroscopeVIS2.0Pro software (Kern) to capture images of 0.26 mm^2^ areas and quantified stomatal density (stomata mm^-2^) and index (proportion of stomata from all epidermal cells) with ImageJ (Schneider *et al*., 2012). Data were analysed with one-way ANOVA followed by a Tukey *post hoc* test, multiple t-tests with Bonferroni correction, or linear regression as indicated in the figure legends. OriginPro 2018 (OriginLab, Massachusetts, USA) was used for statistical analysis, all effects were considered significant at *P*<0.05.

## Acknowledgements

We thank Sten Mander from the University of Tartu Botanical Garden for providing fern spores for *A. capillus-veneris*, *A. polyphyllum*, *C. distans*, *C. farinosa*, *C. lanosa*, *A. scolopendrium*, and *D. sieboldii*. This work was supported by the Estonian Research Council (grants PRG537 and PSG404), and the European Regional Development Fund (Centre of Excellence EcolChange).

## Author contributions

Ü.N. initiated the work with fern spores. H.H. conceived and designed the study, T.W. and H.H. collected data, T.W. and H.H. analysed data, Ü.N. contributed to data analyses. H.H. wrote the manuscript with T.W. and Ü.N.

