## Supplementary Figure 1 for "Species-specific stomatal ABA responses in juvenile ferns grown from spores"

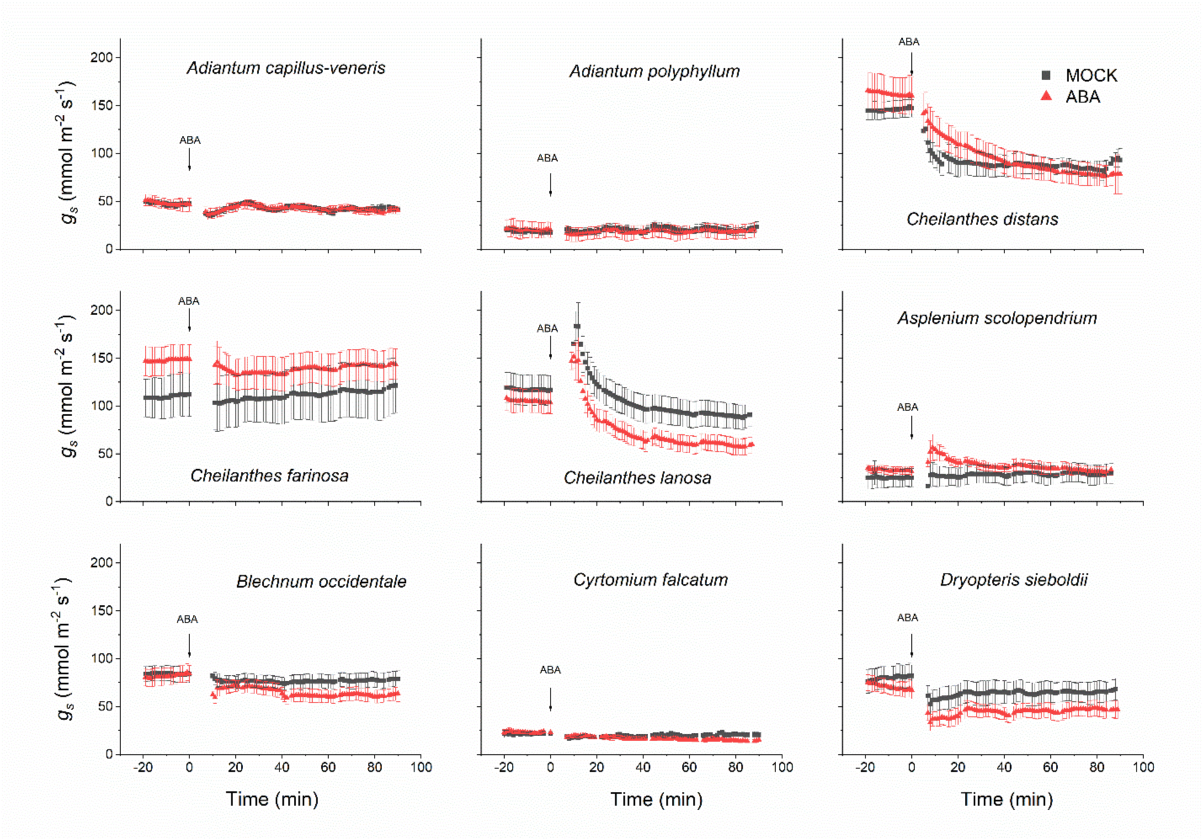


**Supplementary Figure 1. Stomatal ABA responses in juvenile ferns.** Stomatal response in absolute units to treatment with 10 µM ABA or mock-solution at time point 0 in *A. capillus-veneris*, *A. polyphyllum*, *C. distans*, *C. farinosa*, *C. lanosa*, *A. scolopendrium*, *B. occidentale*, *C. falcatum*, and *D. sieboldii*. Mean ± SEM is shown in all panels, n=4-5 plants.
